# Predictive posture stabilization before contact with moving objects: equivalence of smooth pursuit tracking and peripheral vision

**DOI:** 10.1101/2024.04.07.588455

**Authors:** Oindrila Sinha, Taylor Rosenquist, Alyssa Fedorshak, John Kpankpa, Eliza Albenze, Cedrick Bonnet, Matteo Bertucco, Isaac Kurtzer, Tarkeshwar Singh

## Abstract

Postural stabilization is essential to effectively interact with our environment. Humans preemptively adjust their posture to counteract impending disturbances, such as those encountered during interactions with moving objects, a phenomenon known as anticipatory postural adjustments (APAs). APAs are thought to be influenced by predictive models that incorporate object motion via retinal motion and extra-retinal signals. Building on our previous work that examined APAs in relation to the perceived momentum of moving objects, here we explored the impact of object motion within different visual field sectors on the human capacity to anticipate motion and prepare APAs for contact between virtual moving objects and the limb. Participants interacted with objects moving towards them under different gaze conditions. In one condition, participants fixated on either a central point (central fixation) or left-right of the moving object (peripheral fixation), while in another, they followed the moving object with smooth pursuit eye movements (SPEM). We found that APAs had the smallest magnitude in the central fixation condition and that no notable differences in APAs were apparent between the SPEM and peripheral fixation conditions. This suggests that the visual system can accurately perceive motion of objects in peripheral vision for posture stabilization. Using Bayesian Model Averaging, we also evaluated the contribution of different gaze variables, such as eye velocity and gain (ratio of eye and object velocity) and showed that both eye velocity and gain signals were significant predictors of APAs. Taken together, our study underscores the roles of oculomotor signals in modulation of APAs.

**New and Noteworthy:** We show that the human visuomotor system can detect motion in peripheral vision and make anticipatory adjustments to posture before contact with moving objects, just as effectively as when the eye movement system tracks those objects through smooth pursuit eye movements. These findings pave the way for research into how age-induced changes in spatial vision, eye movements, and motion perception could affect the control of limb movements and postural stability during motion-mediated interactions with objects.

## Introduction

When interacting with physical objects that are moving with respect to the body or are stationary in external space during our self-motion, we must stabilize posture against the interactive forces. Previous studies by Lacquaniti and colleagues showed that healthy adults estimate the momentum of the object and scale anticipatory postural adjustments (APAs) in the upper limb in preparation of impact (1). Furthermore, these studies have shown that humans fine-tune their APAs to handle changes in momentum, regardless of whether these changes are due to variations in mass or velocity. This ability suggests that humans use a predictive model based on visual cues of the object’s motion or sensorimotor memories of object properties to foresee the outcome of physical collisions (2).

In our previous work, we addressed how humans estimate object motion to prepare APAs (3, 4). Humans process motion through signals received directly from the eyes (retinal signals) when neural signals of an object’s image moving across the retina (see Fig. 1A) are processed by neurons sensitive to motion in the brain’s visual cortex and other motion-processing areas. Additionally, commands for smooth pursuit eye movements (SPEM)—movements that allow our eyes to smoothly follow moving objects—provide crucial extraretinal information (Fig. 1A). These motor commands help the brain understand the movement of objects by keeping the object in sharp focus on the fovea (5, 6), the part of the eye where vision is sharpest, and likely by providing an efference copy to an internal model of object dynamics (7). This dual approach of using both retinal and extraretinal cues enables accurate motion prediction and efficient interaction with moving objects.

**Figure 1:**
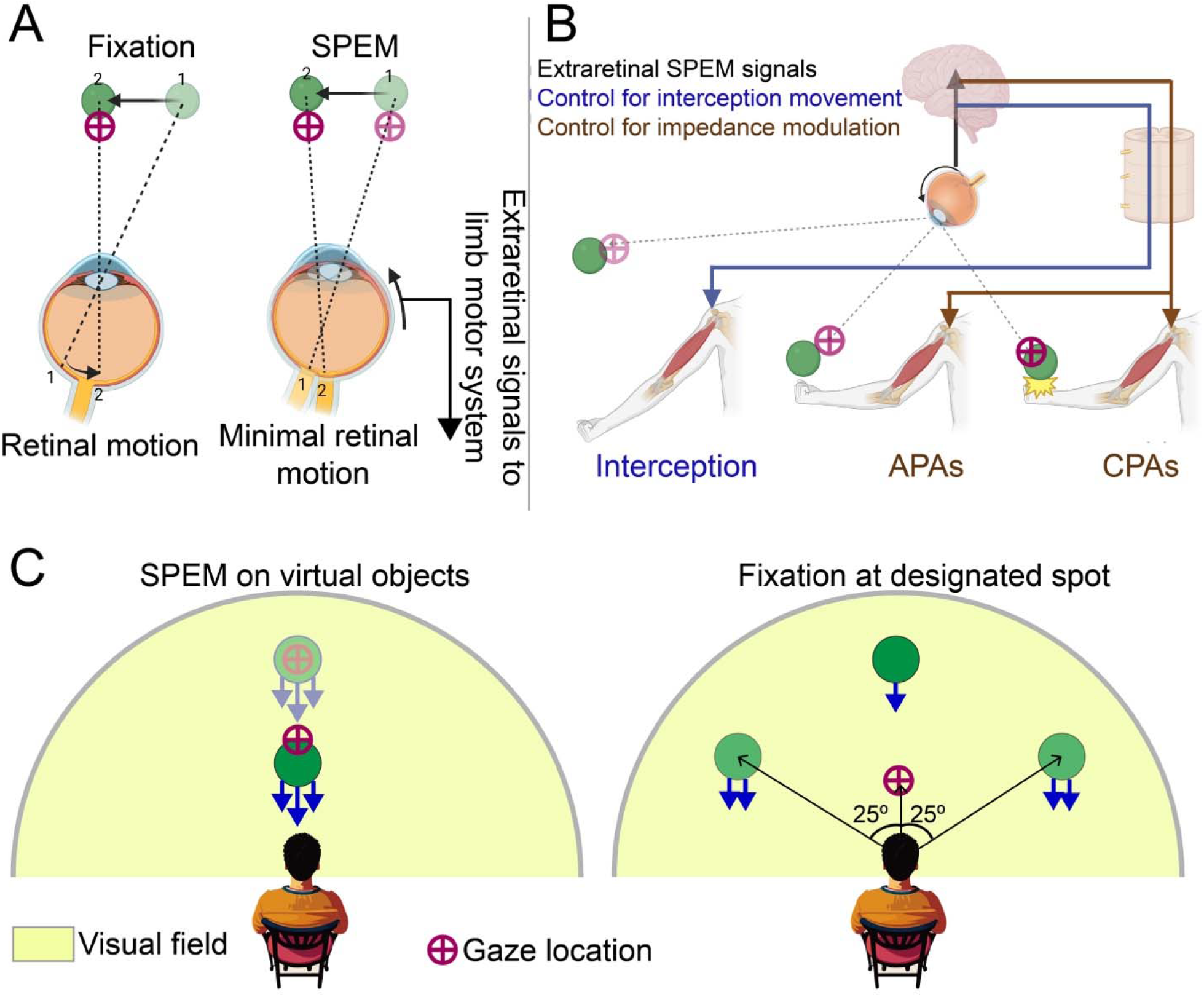
Conceptual framework. A) Retinal motion is produced when eyes are stationary and smooth pursuit eye movements (SPEM_ keep the image on the fovea relatively stationary but produce extraretinal signals for motion perception and motor control. B) Conceptual model for the hypothesis that an efference copy of the extraretinal SPEM signals (black arrow) serve as an input to the limb motor system for control of interception movements (blue arrow) and anticipatory postural adjustments (APAs) prior to contact (brown arrow) and compensatory postural adjustments (CPAs) (brown arrow) during contact with moving objects. C) We have previously shown that the amplitude of APAs is greater when participants track objects with SPEM, (illustrated in the left panel) compared to when they fixate on a point and an object moves along the mid-sagittal plane (dark green circle in the right panel). We attributed this discrepancy to a reduced estimation of object velocity during fixation, as symbolized by a single arrow during fixation versus three arrows during SPEM (left panel). Here, we investigated whether this impaired perception of motion also affects other areas of the peripheral visual field, resulting in decreased APA amplitudes. In this study, the virtual object was moved at a 25° eccentricity within both the left and right visual fields. We posited that, relative to SPEM, object motion in the peripheral visual field would lead to compromised APAs due to diminished velocity estimation, though this effect would be less pronounced (indicated by two arrows) than that observed during fixation along the mid-sagittal plane.

A body of research suggests that an efference copy of extraretinal oculomotor signals is shared with the limb motor system, enhancing the precision of actions aimed at intercepting moving objects (8). Individuals perform interception tasks with greater accuracy when they track moving targets using SPEM (see Fig. 1B) rather than by fixing their gaze on a single point (9–11). Fixed gaze leads to an overestimation of a moving target’s velocity, a phenomenon documented since the works of Aubert in 1886 and Fleischl in 1882 (12–14). This misperception, termed the Aubert-Fleischl phenomenon, leads to an overestimation of the target’s velocity, causing interception movements to be inaccurately aimed ahead of the target (10). Yet, our recent study (4) indicates that subjects instructed to stop a virtually moving object by applying a force pulse will exert less force with a fixed gaze, suggesting an underestimation rather than an overestimation of the object’s velocity. Our recent findings challenge the idea that people overestimate motion when their gaze is fixed and indicate that factors other than motion overestimation might explain why individuals tend to overshoot the target during interception tasks yet decrease the magnitude of their APAs when interacting with moving objects.

When gaze is fixed, retinal motion signals are processed with varying degrees of accuracy in different parts of the human visual field. For example, peripheral vision has superior detection of motion, especially for horizontal movement (15–17). But slower speed perception in peripheral vision causes delays in the perception of motion onset (18). In previous motor control studies, objects moved across the viewer’s field of vision either in the left (11) or the right visual field (10) with respect to the fixed gaze location. In our previous study (4), we had objects moving downwards in the upper part of the visual field. Additionally, the uneven distribution of cones and rods on the retina influences spatial resolution across the visual field, with a more pronounced decline in vertical than horizontal directions. This form of vertical meridian asymmetry is termed a “performance field” (19, 20). Together, this raises the possibility that asymmetrical motion processing in the visual field may impacts APAs for interactions with moving objects.

Here, we first examined APA timing and magnitude when the approaching object is tracked with smooth pursuit eye movements or during fixation such that the object moves within different areas in the field of vision. We hypothesized that signals from SPEM would offer a more accurate estimate of the object’s motion compared to motion signals in the peripheral vision. Based on this, we predicted that the strongest APAs would occur when objects are being actively tracked with SPEM, followed by objects moving in the left and right visual fields, and least strong for objects moving in the upper visual field (Fig. 1C).

To test our hypothesis, we also investigated which gaze signal most accurately predicts APAs. In oculomotor control studies, SPEM gains or gaze gains (the ratio of eye to target velocity) are often used to measure tracking accuracy. Typical gain values range from 0.3-1.1 and are lower for targets moving at high velocities (21, 22). In eye-hand coordination tasks, higher target velocities demand faster eye and interception movements and necessitate appropriate APA scaling to absorb the object’s momentum during contact. Therefore, eye movement velocity may also influence APA modulation. The correlation analysis between gaze signals and APAs was conducted only for conditions where objects were actively tracked using SPEM. In fixation conditions, gaze velocity was close to zero, making this analysis infeasible. We tested SPEM gain (the ratio of eye to object velocity) (21, 23–25), unnormalized eye velocity, retinal slip, and spatial discrepancy of the object and gaze as potential predictors of APAs.

In brief, we found that during fixation, the accuracy of APAs based on the perception of object motion in the peripheral visual field is comparable to that during smooth pursuit eye movements (SPEM). In contrast, when an object moves along the mid-sagittal plane of the visual field, its motion is likely perceived as slower, leading to reduced APA indices. We also found that gaze gains and unnormalized gaze velocity signals were the best predictors of APAs.

## Methods

### Participants

Sixteen right-hand dominant individuals (mean age: 21.3 ± 1.5 years; 9 males and 7 females) participated in the study. All the participants were screened for any history of neurological conditions or musculoskeletal injuries in the upper limb. Before participating, each participant provided written informed consent and was compensated for their time ($10/h). All study protocols were approved by the Institutional Review Board of the Pennsylvania State University.

### Apparatus and stimuli

Participants performed the experimental tasks using a Kinarm Endpoint robot (Kinarm, ON, Canada) equipped with a force transducer within the handle. This Kinarm setup was coupled with an SR EyeLink 1000 Remote eye-tracking system (SR Research, ON, Canada). To interact with virtual stimuli projected from a monitor onto a semi-transparent mirror above the workspace, obstructing direct vision of the hand, participants grasped a robotic manipulandum with their right hand. Their head was tilted approximately 30° forward during the experiments. Visual stimuli, including the cursor indicating the current hand location, were displayed at 120 Hz on a virtual reality display (VPixx Technologies, QC, Canada). Additionally, electromyographic (EMG) data were collected from two arm muscles, the biceps brachii and triceps brachii, using wireless surface electrodes (Trigno, Delsys Inc, MA, USA). Hand kinematics and kinetics were recorded at 1,000 Hz. The virtual moving stimuli was generated using a Gabor-patch generator (https://www.cogsci.nl/gaborgenerator), with properties identical to those in our previous studies (3, 4).

The monocular eye-tracking system was mounted ∼80 cm in front of the participant’s face. For our data collection we maintained a dark atmosphere and used the built-in 13-point calibration provided by KINARM. Eye movements were recorded with a maximum sampling frequency of 500 Hz and accuracy of 0.25-0.5°. Slight left and right head movements were permitted, with the participants keeping their forehead on a resting pad placed on the KINARM. The system records left eye movement along the x and y axis of the KINARM’s visual surface. Data loss with our device was approximately (2.6 ± 0.9%) (26).

### Experimental task

We employed a task called the Mechanical Stopping of a Virtual Projectile (MSTOP) (3, 4), where participants had to stop a virtual object moving towards them at a steady velocity. This task was designed to simulate the physical interaction with an object by assigning it a virtual mass. The collision effect was created by calculating the object’s momentum, which was the product of its virtual mass and velocity (Eq. 1a). When the object encountered the participant’s hand, this momentum was transformed into a force impulse (Eq. 1a), delivered by a robot as a trapezoidal force profile over a set period (90 milliseconds, with rise and fall times of 10 milliseconds each, Fig. 2A). Participants were required to generate a counterforce that fell within a specific error range (±15%) to successfully stop the object (Eqs. 1b-c). The difference between the hand-applied and robot-applied impulses (as shown in Eq. 1c) needed to be within 85% to 115% of the ideal value. The goal was to achieve a ΔImpulse of zero for a perfect stop. After each attempt, participants were given both qualitative and quantitative feedback on their performance, indicating how close they were to achieving the ideal ΔImpulse (see Fig. 2B).

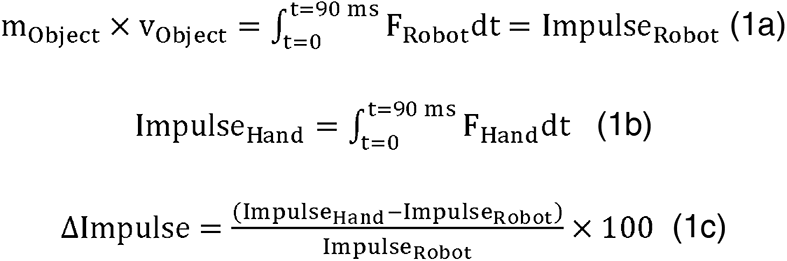

**Figure 2:**
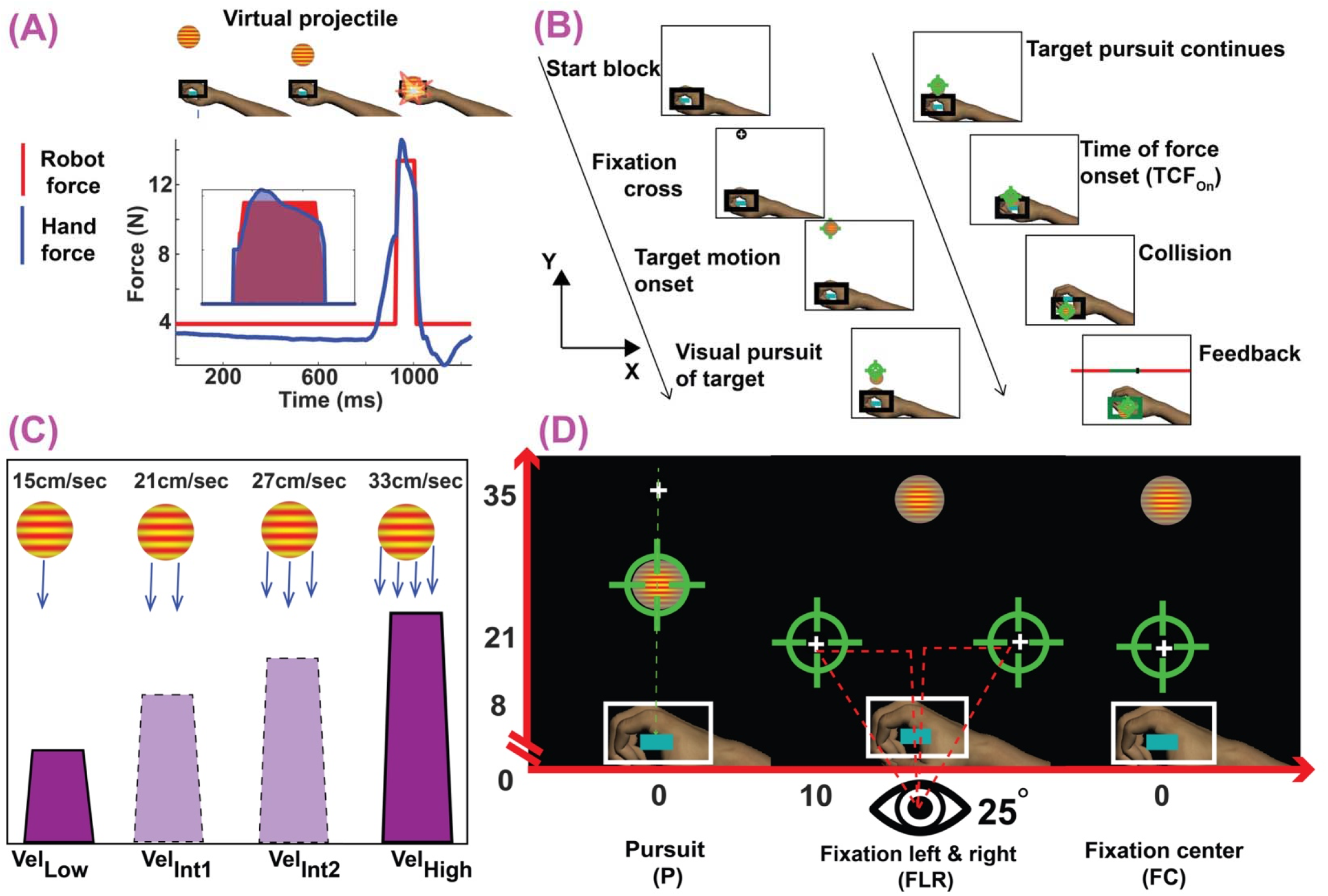
Experimental framework. A) MSTOP paradigm. Participants were required to stop a virtual object moving towards them. When the object reached the hand, the robot applied a force impulse on the hand based on Eq. 1. Participants were required to apply a force on the manipulandum (blue curve) to match the robot impulse (area under the red curve). The blue area in the inset shows the hand force signals that were integrated to calculate Impulse_Hand_ in Eq. 1b. B) Trial flow. At the beginning of each trial, participants were instructed to move a cursor representing the veridical position of the hand (blue rectangle) to a start position (black outlined rectangle). Then a fixation cross appeared. Participants had to foveate the cross. After the cross disappeared (for pursuit gaze condition), the object appeared and moved toward the participant. For the fixation center (FC) and fixation left-right (FLR) gaze conditions, the fixation cross stayed on, and the participant had to foveate the fixation cross for the entirety of the trial. The x and y-axes of the workspace are also shown. C) Momentum conditions. The four-momentum conditions were created by varying the velocity of the moving objects (15cm/sec – Vel_Low_, 21cm/sec - Vel_Int1_, 27cm/sec – Vel_Int2_, 33cm/sec - Vel_High_). D) Gaze conditions. Participants performed six blocks of SPEM (P), six blocks when gaze was fixated either to the left or right (FLR) of the moving object, and 6 blocks of fixation in the center (FC). The order of the blocks was randomized between participants.

Each trial began with the participants positioning a cursor, which reflected their real hand location, to a designated start point. Simultaneously, a constant background force of 3 Newtons (N) was applied toward the body to minimize premature hand movements before the introduction of the moving object. After setting the hand in position, a fixation cross was displayed (see Fig. 1B), requiring participants to fixate on it for 800 milliseconds while keeping their hand steady within the designated start point. Following the removal of the fixation cross, within a timeframe of 250 to 750 milliseconds, the virtual object was introduced and started moving towards the participant.

### Experimental conditions

We designed four different mechanical scenarios by altering the velocity of the object, while keeping its virtual mass constant at 5.33 kg and its radius at 1.5 cm. The conditions varied in terms of the object’s momentum, which was adjusted through changes in velocity alone. The condition with the lowest velocity, referred to as Vel_Low_, had the object moving at 15 cm/s. The highest velocity condition, named Vel_High_, involved the object moving at 33 cm/s. Additionally, to test our hypothesis, we included two intermediate velocity conditions, Vel_Int1_ and Vel_Int2_, with the object’s velocities set at 21 cm/s and 27 cm/s, respectively (see Fig. 2C).

### Study design

The study was carried out over two days.

#### Day 1

The study was carried out over two days. On the first day, participants were familiarized with the slowest (Vel_Low_) and fastest (Vel_High_) object speeds through practice. They completed four blocks of 30 trials for each speed, focusing on tracking the objects using smooth pursuit eye movements (SPEMs). This tracking condition is referred to as the Pursuit (P) condition (see Fig. 2D). It is important to note that SPEM, as used here, encompasses both smooth pursuit and vergence eye movements. While smooth pursuit movements are conjugate (both eyes move in the same direction) and vergence movements are disconjugate (eyes move in opposite directions), they have been traditionally studied separately due to their differing control mechanisms. However, research indicates considerable overlap in the brain regions controlling these movements, suggesting a shared regulatory mechanism (27–29) which is consistent with the fact that these eye movements typically occur together during interactions with objects moving towards the body or when the body moves towards stationary objects. For simplicity, we will use the term SPEM to collectively describe both smooth pursuit and vergence eye movements in our study.

#### Day 2

On the second day of the study, participants began with a set of 16 EMG normalization trials. In these trials, when the participant’s hand contacted a virtual target, they encountered a 10 N force directed either towards them (in 8 trials) or away from them (in 8 trials). They were required to maintain their arm’s position against this force, and the collected data was later used for normalization purposes.

After completing the normalization trials, participants performed 18 blocks of trials using the Mechanical Stopping of a Virtual Projectile (MSTOP) paradigm, divided equally among three different gaze conditions: Pursuit, Fixation center, and Fixation left-right, with six blocks for each condition. The order of these gaze conditions was randomized for each participant. In the fixation conditions, the visual fixation point was positioned either five cm from the starting point in the Y direction along the midline of the body for the Fixation center (FC) condition, or 25° to the left or right of the central line for the Fixation left-right (FLR) condition. Participants were instructed to keep their gaze fixed on the cross for the duration of these trials, and the experimenter monitored their gaze to ensure adherence to the instructions. Each block consisted of 30 trials, including 12 trials each at the Vel_Low_ and Vel_High_ velocities, and 3 trials each at the intermediate velocities Vel_Int1_ and Vel_Int2_, with the order of trials randomized within each block. The idea for having only 10% of trials each of Vel_Int1_ and Vel_Int2_ (intermediate velocity conditions) in a block was to probe how participants use oculomotor and motion signals to prepare APAs in less-practiced conditions. If participants used learned forward models to prepare APAs based on the familiar low and high velocity conditions, we would expect APAs to be similar between Vel_Low_ and Vel_Int1_, as well as between Vel_Int2_ and Vel_High_.

### Data recording and analyses

#### Kinetic and kinematic analysis

The study analyzed hand movement using both kinematic (motion) and kinetic (force) data. To prepare the data, we applied a low-pass filter at 50 Hz to kinetic variables and 15 Hz to kinematic signals, using a double-pass, zero-lag, 3rd order filter for both. Before contact between the hand and the virtual object, participants increased their hand force and moved slightly towards the object, usually reaching maximum force just before contact. This maximum force at contact is termed anticipatory peak force (AF_Peak_, see Fig. 3A, left panel). We then analyzed the force data backwards in time to identify when the force fell below 5% of AF_Peak_ and located the nearest minimum force or inflection point. The earliest of these points marked the hand force onset (AF_On_). By fitting a first-order polynomial between AF_On_ and AF_Peak_, we calculated the slope to quantify the rate of force development (AF_Rate_). At AF_On_, we measured the distance between the hand and the object and divided it by the object’s speed, to determine the time of force onset from contact (TCF_On_).

**Figure 3:**
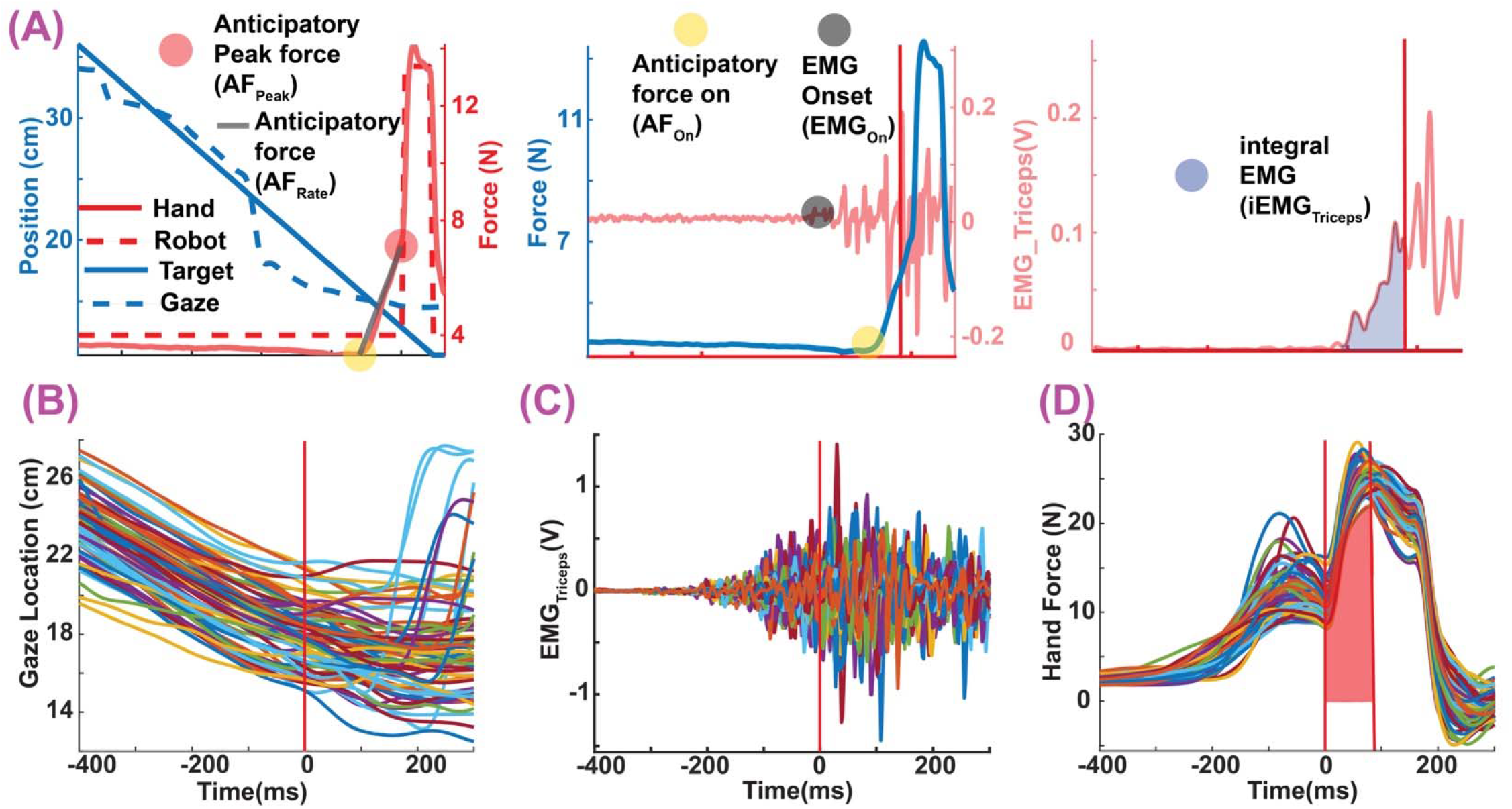
Parameters of interest and exemplar trials. A) The first panel shows object (blue solid line) and gaze (blue dashed line) locations (along the y-axis of the workspace) on the left y-axis and the robot (red dashed line) and hand (red solid line) force on the right y-axis for one trial. Rate of force development (AF_Rate_), and anticipatory peak force (AF_Peak_) are also shown. The middle panel shows the hand force (blue solid line) on the left y-axis, the onset of muscle activity in the triceps (EMG_On_) and hand force (AF_On_). The last panel shows how the integral of triceps EMG (iEMG) was calculated based on the area under the EMG signal between EMG_ON_ and the collision point (red line). B–D) example trials showing the gaze location along the y-axis of the workspace, EMG of Triceps Brachii, and hand force in the y direction of one block of Vel_High_ condition for one exemplar participant. The red area in D) shows the 90 ms over which hand force signals were integrated (Eq. 1b).

#### EMG analysis

To analyze the electromyographic signals, we first applied a 5 Hz high-pass filter, followed by signal rectification, and a 50 Hz low-pass filter. For normalization trials, we subtracted rectified baseline signals prior to background load onset and then calculated root mean square (RMS) on the processed signals to derive a single RMS value for each muscle. We normalized the EMG signals during MSTOP trials by dividing them by the RMS values. To determine the onset of EMG activity (EMG_On_, Fig. 3A middle panel), we used the Teager–Kaiser energy operator setting the detection threshold at four standard deviations above the mean (30). Each trial underwent visual inspection to ensure the detected EMG onset was accurate and meaningful. Finally, we integrated the EMG activity for each condition and participant using trapezoidal numerical integration (via the “trapz” function in MATLAB). This integration occurred between the onset of EMG activity (EMG_On_) and time for AF_Peak_ (Fig. 3A, right panel). The signal-to-noise ratio in the biceps signal was low for most participants, leading to its exclusion from further analysis.

#### Gaze data analysis

Gaze data were preprocessed and analyzed as detailed in our prior work (4, 31), involving low-pass filtering point-of-regard (POR) data at 15 Hz, converting it to spherical coordinates, and calculating ocular kinematics. Angular gaze speed was refined using a Savitzky-Golay filter (6th-order, 27-frame window). We classified gaze events—saccades, fixations, and smooth pursuits—using a threshold-based algorithm (31). This method effectively distinguished between various gaze behaviors within trials, including smooth pursuits, saccades, and fixation on a central point during fixation trials. Exemplar gaze, EMG, and hand force data for one participant are shown in Figures 3B-D.

### Bayesian model averaging to determine the gaze variables that predict APAs

We compared multiple predictor variables of gaze and object motion and position that could predict the anticipatory postural adjustments (APAs) prior to collision between the hand and the virtual objects. Specifically, we compared gaze velocity (*Ġ*), retinal slip (*Ġ*−*Ṫ*), gaze gain (*Ġ*/*Ṫ*) and object-gaze distance (*G-T*). We integrated these values between the time of object onset and increase in hand force above baseline levels (AF_on_) and used them to predict AF_Rate_ and AF_Peak_. In between the instance of object onset and AF_On_, we ignored the time windows when the gaze was not in smooth pursuit of the object (see Fig. 4).

**Figure 4:**
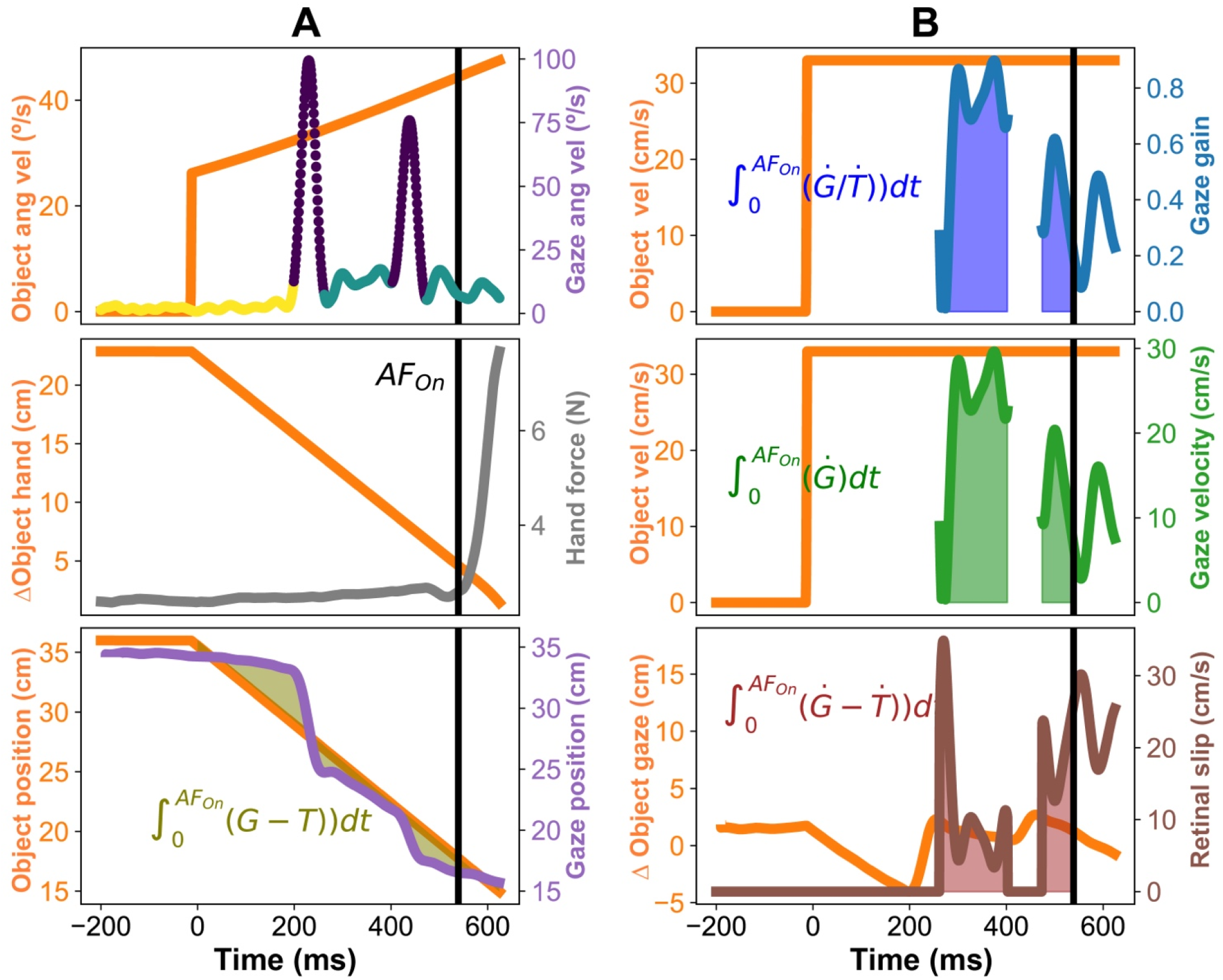
Gaze variables to predict anticipatory postural adjustments (APAs). A) the top panel shows object angular velocity (left y-axis) and gaze ang velocity (right y-axis) with gaze events highlighted: fixation (yellow), saccades (black dots), and smooth pursuits (green). Object angular velocity increases with respect to the eye because the object moves towards the body with a steady Cartesian velocity. Middle panel shows the distance between object and hand as a function of time and the hand force. The onset of hand force (AF_On_) is also highlighted. The gaze variables were only integrated between object onset and AF_On_. Lower panel shows the integrated object-gaze (G-T) distance. B) The three other variables that were tested: integrated gaze gain(*Ġ*/*Ṫ*) in the top panel, integrated gaze velocity (*Ġ*) in the middle panel, and integrated retinal slip(*Ġ*−*Ṫ*) in the bottom panel.

SPEM gain, the ratio of eye to object velocity, is a widely recognized measure of oculomotor performance (21, 23–25). Ideally, the gain approaches 1, indicating precise tracking. In practice, gain values also vary with object velocity (25) and experimental conditions (32), and are notably lower in older adults (21, 33, 34) and various clinical groups (35, 36), suggesting its potential to influence APAs by encoding normalized eye and object velocities. SPEM gain also has inherent limitations since SPEM can reach velocities up to 100°/s, making it difficult to differentiate between objects moving at different velocities. An alternative measure, the unnormalized eye velocity, could better distinguish between objects approaching at different velocities. Another potential signal is retinal slip (37, 38), the difference between object and eye velocity, which is thought to influence the execution of catch-up saccades during SPEM. They are hypothesized to play a role in the programming of catch-up saccades during SPEM (37). To assess the contributions of these signals to APA modulation, we applied Bayesian Model Averaging (BMA) to evaluate their relative contributions. The set of four models is shown below in Equation 2.

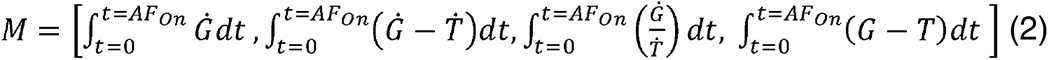

Inferring object motion likely involves evidence accumulation over time, as demonstrated in many random dot kinematogram studies. Therefore, we integrated the oculomotor variables from the onset of object motion to the initiation of hand force (Fig. 3A). We assumed that the evidence accumulation process would be complete before the participant increased hand force above baseline levels.

We implemented Bayesian model averaging for linear regression to compare our predictors (39, 40). This approach allowed us to evaluate 16 models (i.e., 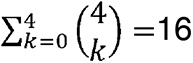, k=0 indicates model with constant term only) across four predictors 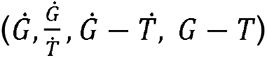 by computing all possible combinations. Initially, we set the cumulative likelihood for all models to zero. By iterating through each predictor variable, we calculated the linear regression and determined the model’s likelihood using the BIC index. The essence of model averaging lies in calculating the inclusion probability of each predictor variable, which is the ratio of the sum of likelihoods for all models including that variable to the total likelihood. BMA circumvents the binary approach of classical hypothesis testing, where models are either entirely accepted or rejected, and is resilient against errors in model specification. The output of the analysis is a probability value that shows how likely is this variable to be included into the 2^4^ (16) combinations of models. A value of 1 would suggest that any model that does not include this term is extremely unlikely.

### Statistics

On Day 1, participants learned the new task, and data from this day was not analyzed further. On Day 2, our objectives were to: a) examine the impact of the three gaze conditions (pursuit, fixation left right, and fixation center) and object speed on our experimental variables; and b) determine which gaze variables predict APAs. We began with testing for normality using the “normaltest” function from the Pingouin Python package. We then assessed homoscedasticity, or the equality of variances across groups, using the Bartlett test. Finally, we checked for sphericity with the “sphericity” function from the Pingouin package. Any deviations from these assumptions were addressed through data transformations. Following these preparatory steps, we conducted a repeated measures ANOVA (RM-ANOVA) to understand the effects of Gaze Condition (Pursuit, Fixation center, and Fixation left-right as levels) and Object Velocity (Vel_Low_, Vel_Int1_, Vel_Int2_, and Vel_High_ as levels) on our dependent variables. The analysis considered gaze and speed conditions as within-subject factors, with participants treated as random effects. We calculated effect sizes (generalized η^2^) to evaluate the strength of the relationships. Alpha level was set to 0.05. For detailed comparisons, post hoc pairwise tests with Holm correction were performed to quantify the differences between conditions.

We then tested if our dependent variables (TCF_On_, iEMG_Triceps_, and AF_Peak_) changed differently with increase in object velocity across the three gaze conditions (Pursuit, Fixation center, and Fixation left-right). Did these variables increase at the same rate, regardless of the gaze condition, as the object moved faster? To determine this, we examined the slopes of a multivariate linear regression relating the variables to object velocity. This method gave more credence to measurements that were more reliable (i.e., smaller variance) for different velocities. We then compared these slopes across the three gaze conditions to see if they were consistent. The comparison involved calculating a test-statistic that represents the difference in slopes between each condition and the average slope across all conditions. Then to test if these differences were significant, we randomly shuffled the data 20,000 times randomly, assigning the data points to different gaze conditions each time but keeping their object velocity grouping the same. After each shuffle, we recalculated the test-statistic. Finally, the p-value was then calculated as the proportion of permuted test statistics that were greater than or equal to the observed test statistic. We set a standard (α = 0.05) for significance, meaning if the proportion was below this threshold, it indicated that the slopes were not the same across all gaze conditions, suggesting that the variables’ behavior depended on the gaze condition at different object speeds. If the p-value was found to be less than the significance level (α), subsequent post-hoc permutation tests were conducted. These tests aimed to identify specific differences in the slopes of gaze conditions. To accomplish this, the methodology of the initial permutation test was applied to pairs of gaze conditions (e.g., FLR vs. P, P vs. FC, and FC vs. FLR). For each pair, p-values were calculated to determine statistical differences between their slopes.

Statistical comparison of correlations between different sets of variables was conducted using the Dunn and Clark’s z for dependent groups with overlapping variables as implemented in cocor package in R (41). As before, significance levels were set to α = 0.05.

## Results

The participants in this study learned the task similarly to participants in our previous study (3), i.e., by minimizing the variance of ΔImpulse (Eq. 1c) across blocks. We calculated the standard deviation of ΔImpulse for each block and compared it across blocks. Statistical tests confirmed that the standard deviation of ΔImpulse decreased with practice, showing a main effect of block (p<0.05). Since learning was not the focus here, we did not analyze Day 1 data further.

The first goal of the study was to differentiate how motion signals of moving objects are processed through active pursuit eye movements and passive observation in the peripheral visual field. To that end, we asked the participants to fixate their gaze on a cross while the object moved in different parts of the visual field. Our eye-tracking data shows that we were able to successfully constrain participant’s gaze in the fixation left-right (FLR) and fixation center (FC) conditions (see Fig. 5A, upper panel). The average smooth pursuit duration for the Fixation left-right (0.19 ± 0.06%) and fixation center (3.2 ± 0.74%) was significantly less than during the pursuit blocks (57.5± 1.6%). The RM-ANOVA model confirmed a main effect of gaze condition [F(2,30) = 7.5, P < 0.001, η^2^=0.98]. There was also a main effect of velocity condition [F(3,45) = 7.05, P < 0.001, η^2^=0.82] suggesting that as object speed increased, the proportion of time the gaze was engaged in SPEM decreased and more catch-up saccades occurred.

**Figure 5:**
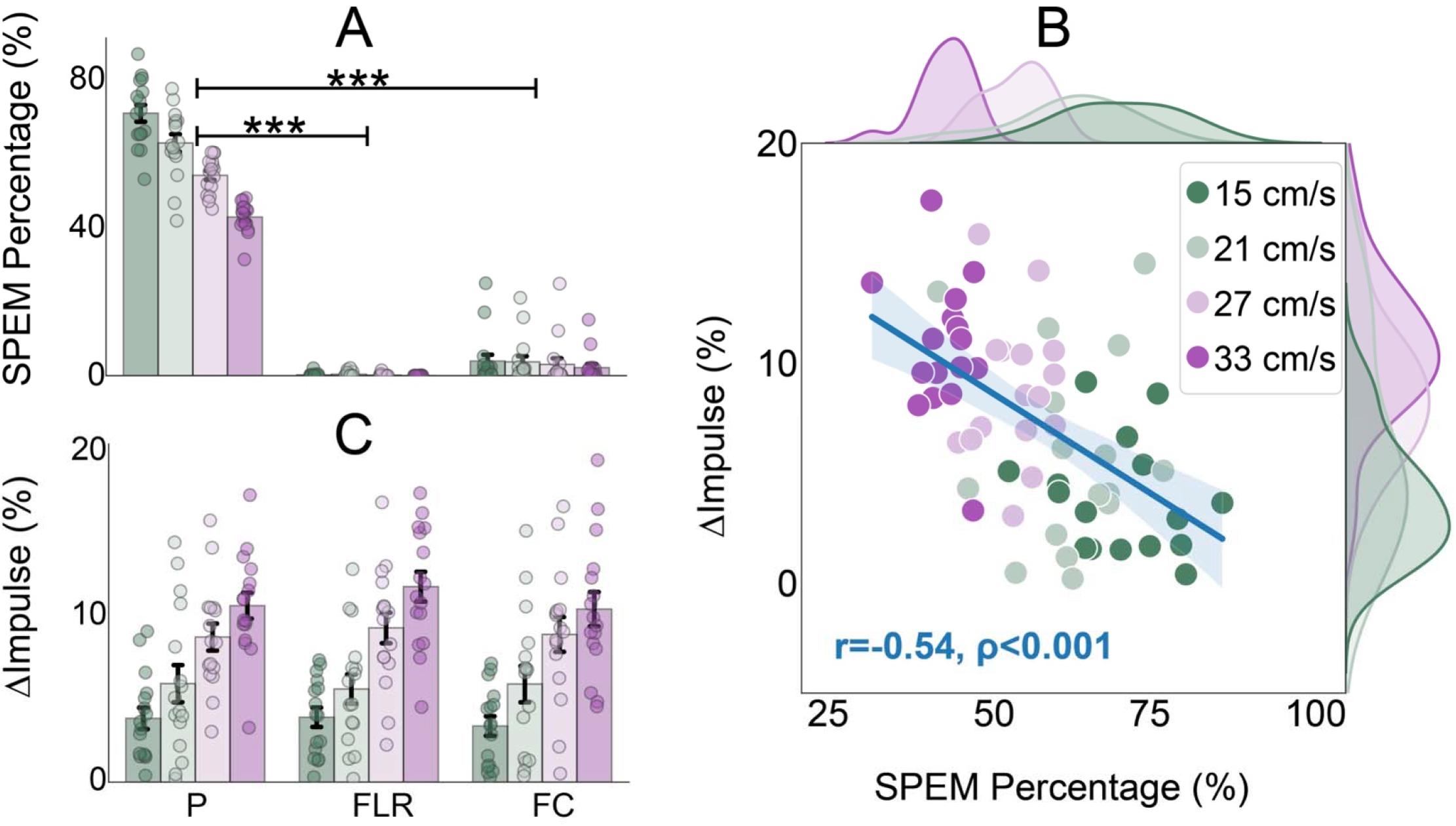
Pursuit eye movements are not causally associated with force control (ΔImpulse, refer to Equation 1c) during contact. A) During the Pursuit condition (P), faster object velocities reduce the percentage of time the gaze is engaged in smooth pursuit eye movements (SPEM), and gaze remained within controlled limits in both fixation scenarios (Fixation Left-Right [FLR] and Fixation Center [FC]). There were significant differences in the percentage of time gaze was engaged in pursuit between the P and FLR and P and FC conditions (*** indicates p<0.001 between different gaze conditions). B) There was a negative correlation between ΔImpulse and the percentage of time spent in SPEM (%SPEM). This inverse relationship suggests that performance deteriorated when the gaze followed the moving objects in SPEM for reduced intervals. C) Imposing restrictions on gaze in the fixation conditions significantly reduced the duration of SPEM across all four velocity settings. The lower panel indicates that the performance variable (ΔImpulse) was similar across the three gaze conditions, but escalated with increasing object velocity, indicating a decline in performance at higher speeds.

The proportion of time spent in smooth pursuit (SPEM) could serve as a predictor of motor performance, quantified as ΔImpulse. A significant negative correlation (r=-0.53, p<0.001) was observed during the Pursuit condition between the duration of smooth pursuit and ΔImpulse, as illustrated in Figure 5B. This indicates that a greater engagement in SPEM was associated with more accurate matching of the robot impulse to stop the object. ΔImpulse declined with increasing velocities of the object, yet it remained comparably consistent across the three gaze conditions, as depicted in Figure 5C. The RM-ANOVA model revealed a significant main effect of the velocity condition on ΔImpulse [F(3,45) = 41.6, P < 0.001, η^2^=0.74], with no other effects reaching statistical significance. This indicates that while SPEM are associated with better task performance in the SPEM condition, there may be other factors that may contribute to similar performances in the fixation conditions (see Discussion).

### Motion processing in different parts of the visual field have heterogeneous effects on APAs

Based on APA results from our previous study (4), we hypothesized that restricting eye movements to enforce motion processing in different parts of the visual field would have a graded impact on decreasing the magnitude of APAs (AF_Rate_, AF_Peak_, and iEMG_Triceps_) and timing variables (TCF_On_) prior to contact between a virtual projectile and the hand. Embedded within this hypothesis is the assumption that extraretinal signals associated with smooth pursuit eye movements (SPEMs) provide the limb motor system with a superior estimate of object motion. Additionally, we predicted that faster-moving objects would lead to stronger APAs and quicker onset times (TCF_On_).

Gaze conditions had no significant effects on two dependent variables of interest TCF_On_ [F(2, 30) =0.05, P = 0.92, η^2^=0.04, see Fig. 6A] and iEMG_Triceps_ [F(2, 30) =0.76, P = 0.47, η^2^ =0.81, see Fig. 6B]. However, as expected, the main effect of velocity was significant for both TCF_On_ [F(3, 45) =43.58, P <0.001, η^2^=0.74] and iEMG_Triceps_ [F(3, 45) =9.47, P <0.001, η^2^ =0.39]. Interaction effects were not significant for either TCF_On_ [F(6, 90) =0.97, P=0.39, η^2^=0.06 or iEMG_Triceps_ [F(6,90) =0.97, P = 0.46, η^2^ =0.06]. This indicates that faster object velocities led to notable changes in the onset time of limb force and the integrated amplitude of the agonist muscle activity. We have not presented data for EMG_On_ (Fig. 4A) because they were similar to TCF_On_.

**Figure 6:**
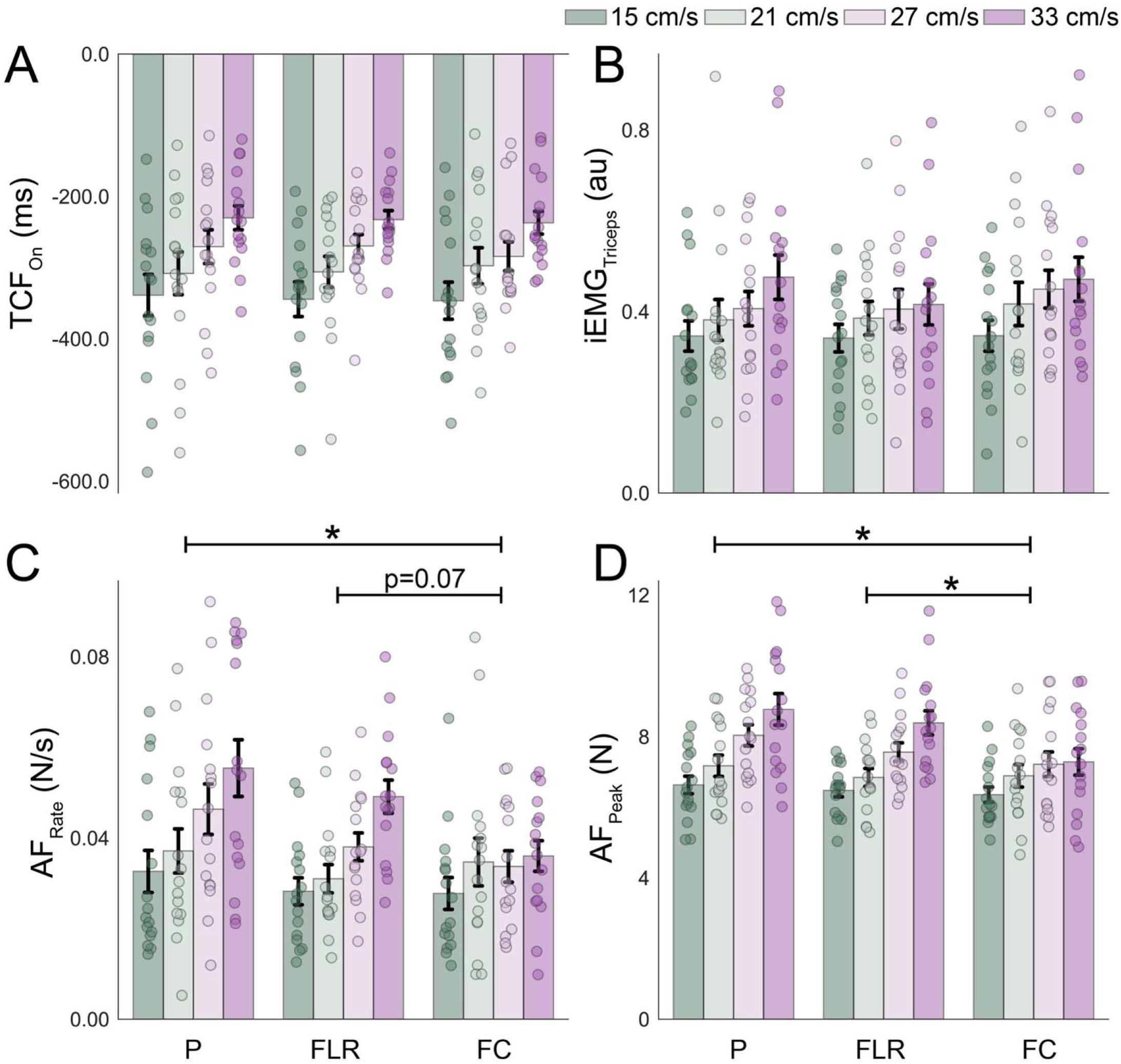
Modulation of anticipatory postural adjustments (APAs). APAs are smaller in magnitude when gaze is fixated and object moves along a mid-sagittal plane. The influence of different gaze conditions on muscle activation and the timing of these activations appears negligible. A) The upper left panel indicates that the timing of force onset (TCF_On_) remains statistically unchanged across different gaze conditions. B) The upper right panel demonstrates that the integrated muscle activity of the triceps muscles (iEMG) is not significantly affected by the type of gaze condition. C and D) The lower panels reveal variations in anticipatory postural adjustments, specifically in the rate of force development (AF_Rate_) and the anticipatory peak force (AF_Peak_), between combinations of gaze conditions (SPEM & fixation center versus fixation left-right & fixation center). However, no significant differences were observed between SPEM and fixation left-right (FLR) conditions. ’*’ indicates significant differences between different levels of the gaze condition (p<0.05). The colors of the bars and the corresponding velocity conditions are denoted at the top right.

When participants fixed their gaze at the center, both AF_Rate_ and AF_Peak_ significantly decreased, indicating a notable impact of central fixation. However, no significant differences were observed between the pursuit condition and fixation in the parafoveal area (near the edges of the gaze field), as illustrated in Figures 6C and D. For AF_Rate_, there was a significant main effect of gaze condition [F(2,30) = 3.6, P < 0.05, η^2^ = 0.19], velocity [F(3,45) = 23.6, P < 0.001, η^2^=0.61] and an interaction effect between the factors [F(6,90) = 3.87, P < 0.01, η^2^=0.21]. For AF_Peak_, there was significant main effects of gaze condition [F(2, 30) = 6.2, P < 0.01, η^2^=0.29], velocity condition [F(3, 45) = 47.2, P < 0.001, η^2^=0.76], and a significant interaction between the two factors [F(6, 90) = 9.67, P < 0.001, η^2^=0.39].

Post hoc analysis revealed significant differences between Pursuit and Fixation center for both AF_Rate_ and AF_Peak_ (P < 0.05), indicating these anticipatory responses are influenced by where participants direct their gaze and whether they pursue the moving object. Significant differences were also found between Fixation left-right and Fixation center for AF_Peak_ (P < 0.05), but not quite for AF_Rate_ (P = 0.07). Comparing Pursuit with Fixation left-right revealed no significant differences for either AF_Peak_ (P = 0.95) or AF_Rate_ (P = 0.81).

### Velocity dependent APA modulation was significantly slower during center fixation

We applied a permutation test, a non-parametric method, to assess differences in the regression slopes across four velocity conditions for each gaze condition, as shown in Figure 7. The analysis revealed that with an escalation in object velocity, the anticipatory variables AF_Peak_ and AF_Rate_ increased at a significantly higher rates in conditions involving Pursuit and Fixation left-right compared to the Fixation center condition (P<0.001, AF_Peak_ illustrated in Figure 7, lower panel). Conversely, the analysis found no statistically significant variances in the slopes related to the timing of muscle activation (TCF_ON_) (P=0.8) or the intensity of muscle activation (iEMG_Triceps_) (P=0.2). Together, these results suggest that there may be differences in estimates of object velocity in the Fixation center condition, that caused a more gradual increase in APA measures as a function of object velocity.

**Figure 7:**
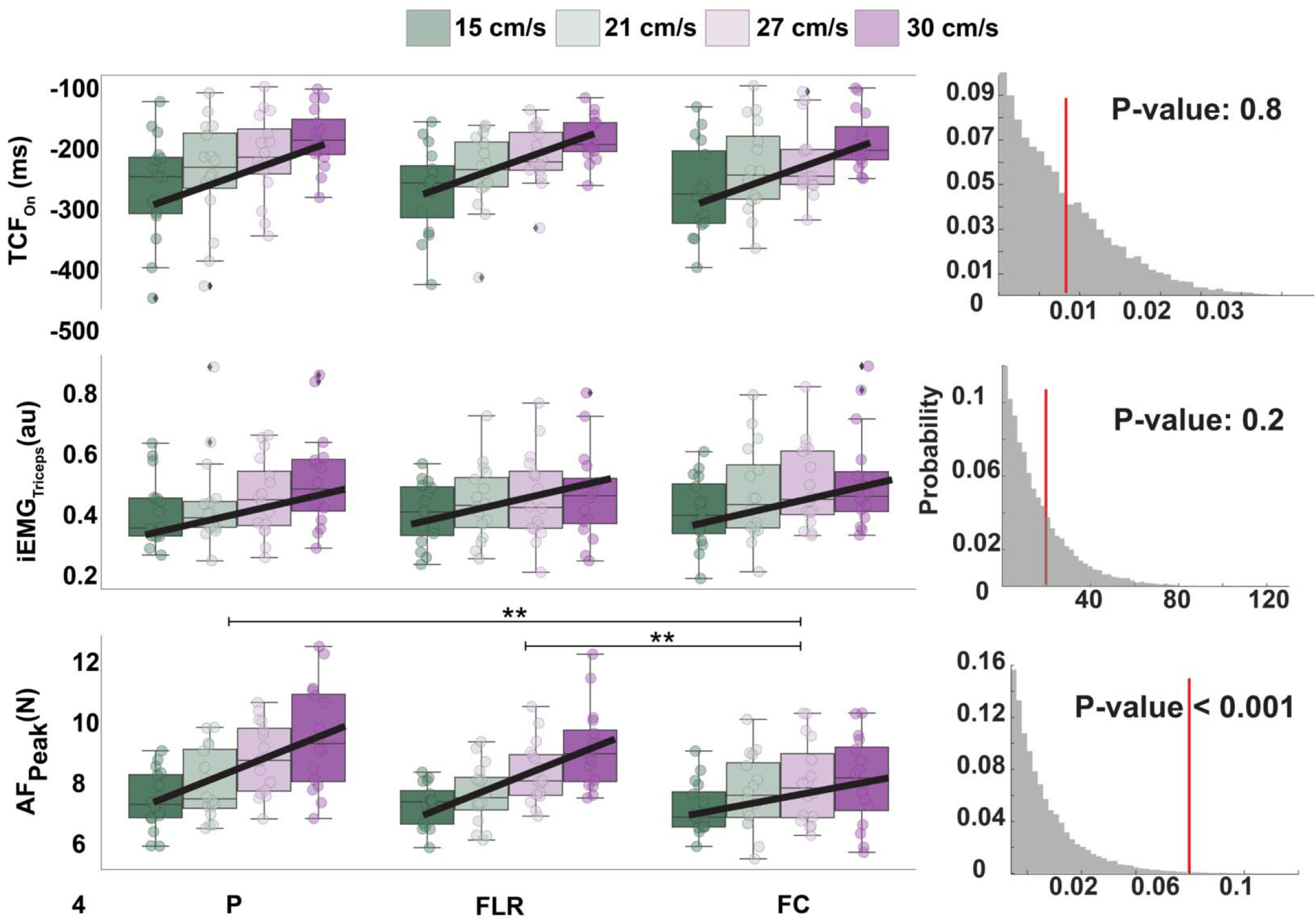
Permutation test analysis exploring the influence of gaze direction on anticipatory posture responses (APAs). Regression models show the relationship between the dependent variables (time of force onset [TCF_On_], integral of triceps muscle activity [iEMG_Triceps_], and anticipatory peak force [AF_Peak_]) and object velocity. The model employs weights derived from the inverse variance within each velocity category to estimate the velocity-dependent rate of change in the dependent variables. Top Panel shows similar slopes for all gaze conditions for TCF_On_, with the permutation test yielding a P-value of 0.8. This suggests that the rates at which TCF_On_ decreases as object velocity increases were similar across the three gaze conditions. Middle Panel shows similar results for iEMG_Triceps_ (p=0.2). Bottom Panel shows the slopes of anticipatory peak force (AF_Peak_) were significantly different across the gaze conditions (p<0.001). Post hoc analyses indicate significant differences in the rate of change between Pursuit and Fixation Center (P<0.01) and between Fixation Left-Right and Fixation Center (P<0.01).

Subsequent analysis utilized post-hoc permutation tests to examine the anticipatory variables, specifically AF_Peak_ and AF_Rate_. These analyses revealed significant differences in the slopes of these variables across various gaze events. Specifically, the slopes differed significantly between the pursuit and central fixation events (P < 0.01), as well as between left-right fixation and central fixation events (P < 0.01), for both AF_Rate_ and AF_Peak_. Conversely, the comparison between pursuit events and left-right fixation events showed no significant difference in slopes for either anticipatory variable, with P-values of 0.47 for AF_Peak_ and 0.38 for AF_Rate_, indicating similar anticipatory responses in these conditions.

### Gaze velocity and gaze gain signals are both similarly effective in predicting APAs

We first calculated Pearson correlation coefficients between either gaze velocity (*Ġ*), retinal slip (*Ġ*−*Ṫ*), gaze gain (*Ġ*/*Ṫ*) or object-gaze distance (*T-G*) as predictor variables and measures of APAs (AF_Rate_ and AF_Peak_) as dependent variables. For both these dependent variables, only gaze velocity (*Ġ*) and gaze gain (*Ġ*/*Ṫ*) were significant predictors. The correlation plots between pairs of variables for AF_Rate_ are shown in Figure 8.

**Figure 8:**
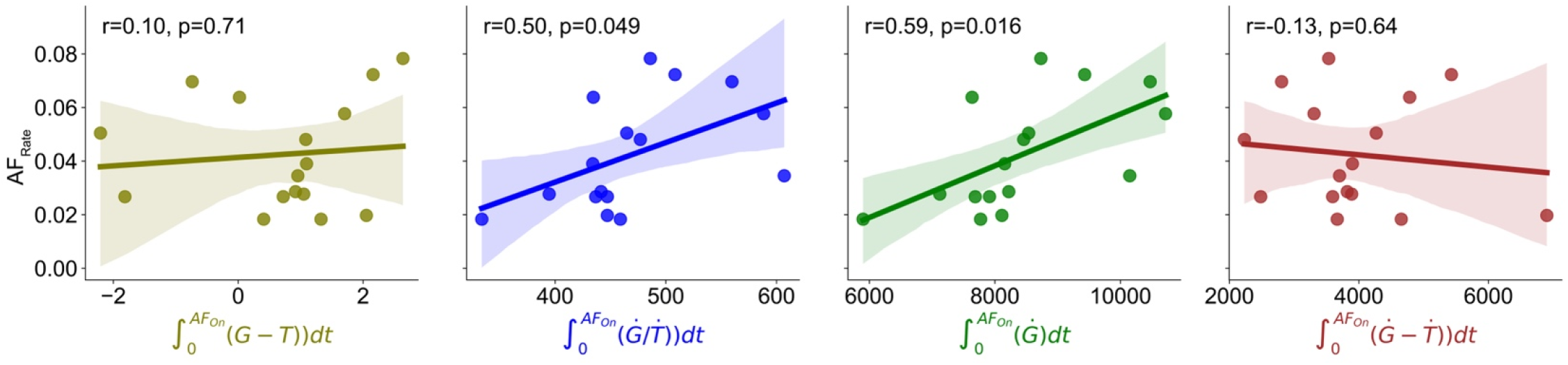
Gaze gain and velocity signals predict APAs. Gaze gain (*Ġ*/*Ṫ*) and gaze velocity (*Ġ*) signals were significantly correlated with APAs, but retinal slip (*Ġ*−*Ṫ*) and gaze-object distance (G-T) were not.

Using the statistical method introduced by (41), we compared the two strongest pairs of correlations - between *Ġ* and AF_Rate_ and *Ġ*/*Ṫ* and AF_Rate_. Our analysis indicated that we could not reject the null hypothesis, which suggests that these two sets of correlations are statistically similar. This outcome was the same when we looked at AF_Peak_. Further, Bayesian Model Averaging (BMA) analysis revealed that gaze velocity (*Ġ*) had the highest likelihood of being an important factor in our model, with gaze gain (*Ġ*/*Ṫ*) ranking next. Notably, these two variables were found to be twice as likely to be included in the model compared to the other two variables we examined (see Table 1). Based on these analyses, we ruled out retinal slip (*Ġ*−*Ṫ*) and object-gaze distance (*T-G*) as predictors of APAs.

**Table 1:** Bayesian model averaging analysis from the regression model for AF_Rate_ (AF_Peak_). This analysis also reveals that gaze velocity (*Ġ* and gaze gain (*Ġ*/*Ṫ*) are the strongest predictors of APAs

| Variable | Probability | Coefficient |
| --- | --- | --- |
| Constant | 0.34 (0.44) | -0.01 (1.45) |
| $\dot{G} - \dot{T}$ | 0.24 (0.25) | -0.003 (-0.26) |
| $\dot{G}/\dot{T}$ | 0.57 (0.47) | -0.08 (2.62) |
| $\dot{G}$ | 0.80 (0.64) | 0.15 (5.36) |
| $T-G$ | 0.20 (0.20) | 0.0004 (-0.01) |

## Discussion

Our findings complement our previous study (4) by revealing a link between motion processing across various parts of the visual field and anticipatory posture stabilization against object collision. Specifically, we show retinal-only object motion signals from the left and right peripheral visual fields enable APAs that are just as effective as those when directly tracking the moving object with SPEM. We also found that gaze velocity and gaze gain are robust and significant predictors of APAs in the upper limb. This outcome supports the premise that signals pertaining to the velocity of eye movements are instrumental in pre-adjusting the posture of the limbs in anticipation of contact.

Different neural pathways connect the cerebellum and brainstem to manage SPEM (28, 42–44) and control posture (45–47). Recent studies have hinted at a functional connection between SPEM and maintaining upright balance (48, 49), which is not unexpected given these findings. Furthermore, damage to the cerebellum – key brain region for coordination leads to slower smooth pursuit and vergence eye movements (50, 51), difficulty adjusting APAs (52), and delayed and altered long-latency reflexes (53, 54). These findings suggest that the cerebellum may be crucial in coordinating SPEM with the mechanisms that stabilize posture.

A key result that confirmed our earlier study is that multiple APA measures – TCF_On_, iEMG_Triceps_, AF_Rate_, and AF_Peak_ – were linked to object speed in all gaze conditions. This consistently supports the idea that the visual system is inherently equipped to detect and process motion with high accuracy and precision, both for perception and for guiding actions. Our findings also indicate that motion can be processed effectively for the purpose of posture stabilization, regardless of whether eye movements were involved. However, this ability to effectively process motion does not extend to the kinematic aspects of limb motor control since gaze fixation degrades the accuracy of movements aimed at moving objects, as discussed in the following section.

### Motion prediction for posture stabilization: The role of smooth pursuit eye movements (SPEM)

It is well established that reaching to peripheral static objects during fixation leads to greater kinematic inaccuracies than when foveating the object (55–57). Similar results have also been shown for moving objects as summarized in three recent reviews (8, 58, 59). Multiple mechanisms enhance kinematic performance when pursuing moving objects with SPEM. First, prediction of future motion of an object is improved when participants track a moving object with SPEM (60, 61); there is also evidence for overestimation of object speed during fixation (10). Second, tracking moving targets that have complex spatial paths with SPEM improves motion prediction and interception performance (62, 63). This is also supported by studies that show that when object motion is fully predictable, pursuit eye movements and hand movements are not correlated (11, 64). Third, SPEM facilitate continuous updates of interception movements under feedback control. And finally, catch-up saccades interfere with motion perception and interception movements (65). This collection of studies underscores that pursuing objects with SPEM can improve motion prediction and performance, while catch-up saccades that often occur when there is retinal slip due to high object speeds, interfere with performance.

The present study presented an object with a fully predictable trajectory and velocity. Increased object speeds led to less accurate motor performance, a result that replicated our earlier findings (4) (see Fig. 5C). It is possible that lower accuracy at higher speeds result from a reduced duration of smooth pursuit eye movements (SPEMs) (Fig. 5B), though the strong negative correlation between SPEM duration and motor error rates (r=-0.53, p<0.001) does not indicate a direct causal link. In fact, our data show that even when eye movements are restricted, as in the Fixation center (FC) and Fixation left-right (FLR) conditions, the quality of motor performance remains largely stable (Fig. 5C).

This finding aligns with results from our previous study (4), suggesting that eye movements may contribute to better task performance (ΔImpulse) in smooth pursuit conditions. But in fixations conditions, participants may use alternative strategies to regulate ΔImpulse. Previously, we found that in pursuit conditions, gaze gains were correlated with anticipatory postural adjustments (APAs), and APAs were correlated with ΔImpulse. In contrast, during fixation conditions, only APA magnitudes were reduced, with no effect on ΔImpulse. This suggests a weak task-induced mechanical coupling between APAs and ΔImpulse. Thus, while SPEM modulates APAs, participants can achieve similar task performance (ΔImpulse) during both SPEM and fixation through other strategies. Further exploration is needed, particularly through careful manipulation of contact mechanics (e.g., Exp. 3 in (4)) and potentially adding a limb movement component to the task.

Restricting eye movements negatively impacts posture stabilization before object contact, but only when the object moved directly in front of the observer along the centerline (or mid-sagittal plane) of the body in the visual field (see Fig. 6). When the object moved through the outer left and right areas of the visual field, the degree of postural stabilization was similar to when the eyes followed the moving object with SPEM. This indicates that the mechanisms of processing motion in the peripheral visual areas are strong and perform on par with the motion information provided by the oculomotor system when eye movements smoothly follow moving objects.

This interpretation is reinforced by an analysis of the regression slopes for different object speeds across the three types of gaze conditions (see Fig. 7). Significant differences were only found between tracking movement with SPEM (Pursuit) and looking straight ahead (Fixation center), and between fixating to the left or right (Fixation left-right) and looking straight ahead (Fixation center). So, APAs based on retinal motion were diminished when looking along the object’s path, the outcomes from the other two gaze conditions were comparable.

The analysis of intermediate velocity trials (Vel_Int1_ and Vel_Int2_) supports the hypothesis that the quality of retinal motion signals along the object’s path is diminished and impacts APAs. These trials served as “catch trials” to determine whether processed motion signals (through extraretinal or retinal mechanisms) directly influenced anticipatory postural adjustments (APAs). If participants used these signals to modulate APAs, we would expect a strong velocity-dependent scaling of APAs, even in these less practiced intermediate velocity trials. Conversely, if participants relied on learned forward models based on familiar low and high velocity conditions to prepare APAs, we would expect similar APAs in the intermediate velocity trials. The gentler slope in the FC condition in Figure 7 suggests that, in the absence of clear processed motion signals, participants may rely more on a learned forward model to adjust APAs.

### Anisotropic motion-processing across the visual field has real-world implications for limb motor control

The human visual system exhibits a sophisticated adaptation to environmental complexities through its anatomical and functional differentiation: the fovea, characterized by its high spatial resolution, facilitates the discernment and manipulation of small objects, contrasting with the peripheral visual field, which is optimized for temporal processing. Notably, spatial visual acuity within the peripheral visual field is non-homogeneous; it diminishes significantly with increasing distance from the fovea because of a sharp decline in the number of photoreceptor cones, exhibiting a more pronounced decline along the vertical meridian compared to the horizontal midline (66, 67). Conversely, the capacity for temporal processing within the peripheral visual field enhances with greater eccentricity, indicating that stimuli located further from the foveal center are processed more swiftly (15, 16). The competing interplay between deficient spatial visual acuity and enhanced motion-processing capabilities in peripheral vision presents a compelling research opportunity to explore the significance of motion-processing in limb motor control. This exploration is relevant for both healthy adults and clinical populations with identified deficits in motion-processing.

Faster temporal processing in the periphery is helpful because it facilitates detection of sudden changes and movements and triggers foveation towards or away from stimuli entering the visual field. Motion processing in the far visual field is also very important for maintaining postural balance during quiet stance and locomotion (17, 68, 69). Initial studies on motion processing have elucidated the anisotropic nature of this capability within the visual field. Specifically, motion processing proficiency is enhanced in the far periphery compared to closer peripheral regions, and the detection thresholds for horizontal motion are lower than those for vertical motion (16). Moreover, velocity discrimination, measured as the relative change in velocity (ΔV/V), is found to be equally accurate in the periphery as it is in the central vision (fovea) (70). Additionally, peripheral vision exhibits minimal deterioration in reaction times and the perception of speed for rapidly moving objects, but not for slower moving objects (71). It is likely that the superior ability of peripheral vision to detect motion, especially of fast moving objects towards the body, is because of its access to specialized neural systems underlying biological motion perception (72).

Our results largely align with these studies that have exclusively focused on motion-processing in the human visual system. Our data indicate that the processing of object motion in the peripheral visual field is as efficient as that achieved through smooth pursuit eye movements, in contrast to the comparatively inferior passive processing observed within the central visual field. In our experiment, the visual stimuli traversed a plane orthogonal to the participant’s body axis. However, due to the participants’ heads being inclined at an approximate angle of 30° from vertical, the stimuli, within the Fixation Center (FC) condition, were perceived to travel along the midsagittal plane but at an oblique angle of approximately 60 degrees relative to the head. Similarly, in the Fixation Left-Right (FLR) condition, stimuli were observed at comparable oblique angles within the peripheral vision. Consequently, an inferred implication of our findings suggests that the human visuomotor system may be more adept at stabilizing posture and engaging with horizontally moving objects, irrespective of their appearance in the central or peripheral visual fields, compared to objects descending vertically. Nonetheless, this proposition is preliminary and requires further empirical validation through more rigorously controlled experimental designs that account for variations in head posture.

### Gaze velocity and gain signals may serve similar roles in limb motor control

Eye movement studies typically report SPEM gains (ratio of eye and object velocity) to determine how the oculomotor system modulates tracking of moving objects as the behavioral conditions change (23, 24, 73). In these studies, visual stimuli are presented on a display oriented in a frontoparallel plane while their heads are stabilized using a chin rest and a bite-bar. The stabilization prevents head movements that would otherwise also require suppression of the vestibulo-ocular reflex. Thus, the SPEM gain is a pure measure of control signals sent to the oculomotor nerves based on continuous negative feedback and motion processing.

In human visual tracking, smooth pursuit eye movements (SPEM) are not the sole mechanism employed for following moving objects; significant head movements are also observed. This behavior is prominently observed in scenarios such as tennis spectators, who actively turn their heads to track the ball’s motion across the court. The development of remote eye-tracking technology, which allows for the measurement of eye movements without restricting head motion, has greatly facilitated the study of more naturalistic eye and head movements (74). Utilizing these advancements, researchers have shifted towards analyzing gaze dynamics through the lens of gaze gain—a metric defined as the ratio of gaze velocity to object velocity, providing a more comprehensive understanding of tracking behavior beyond SPEM gains alone (32, 75, 76).

In our investigation, we incorporated the concept of gaze gain, alongside gaze velocity, to examine their correlation with APA measures. Given that gaze gain values are typically confined within a range of 0.3 to 1.1 (21, 22), we surmised that gaze velocity, which increases with object speed, might offer a superior predictive value for APAs. Our analysis extended to comparing the integrated retinal slip and positional discrepancies between the object and gaze, aiming to identify the most effective predictors of APAs. Determining which of these extraretinal signals is the best predictor of APAs is crucial for testing our hypothesis.

Our findings indicate that both gaze gain and gaze velocity signals are equally effective in predicting APAs, with no significant differences in their predictive capabilities (Fig. 8). This observation implies that both gaze gain and gaze velocity signals demonstrate comparable dynamic temporal modulation during the closed-loop phase of pursuit, where the pursuit response is continually adjusted based on feedback from the motion of the object on the retina. This similarity in modulation likely contributes to their equivalent efficacy in predicting postural responses. In essence, our findings suggest that either signal could be utilized as a predictor. However, the question regarding the precise nature of the extraretinal signal accessed by the limb motor system remains open.

## Conclusions

We examined the how different motion processing signals impact anticipatory postural adjustments (APAs). Specifically, we contrast smooth pursuit eye movement (SPEM) with the processing of object motion while the gaze was fixated directly in line with the object’s motion (FC), or laterally to the left and right of this trajectory (FLR). We observed that APA amplitudes were elevated in the pursuit (P) condition relative to FC, corroborating findings from our prior work (4), but showed no significant differences between P and FLR conditions. During SPEM, various signals including gaze velocity, gaze gain, retinal slip, and the persistent discrepancy between object and gaze positions may influence APA modulation. Utilizing both ordinary linear regression analysis and Bayesian model averaging, we evaluated which of these signals most accurately predicted APAs. Our analysis revealed that gaze velocity and gaze gain were equally effective in predicting APA modulation, outperforming both retinal slip and the spatial discrepancy between the object and gaze. Collectively, these findings elucidate the nuanced advantage of SPEM-derived oculomotor signals in providing motion information for limb motor control, while also underscoring the comparative limitations when set against the intrinsic motion processing mechanisms of the visual system.

## Open Science

All the data and analyses codes from the project are publicly available at DOI: 10.17605/OSF.IO/BK7TE (https://osf.io/bk7te/).

## Notes

### Competing Interest Statement

The authors have declared no competing interest.

### Summary of Updates

1) Figures 3,4,5 and 6 updated. 2) New analysis performed for data collected on Day 1. 3) Provided detailed explanation for inclusion of Bayesian Model Averaging.

